# mTOR complex 1 pathway activation in severe keratoconus; the functional implications of GWAS identified loci

**DOI:** 10.1101/244897

**Authors:** Robert PL Wisse, Jonas JW Kuiper, Gijsbert M de Veij Mestdagh, Catharina GK Wichers, Sanne Hiddingh, Nadia Vazirpanah, Saskia M Imhof, Timothy RD Radstake, Jasper CA Broen

**Affiliations:** Department of Ophthalmology, University Medical Center Utrecht, The Netherlands; Ophthalmo-Immunology group, Laboratory of Translational Immunology, Department of Immunology, University Medical Center Utrecht, The Netherlands; Laboratory of Translational Immunology, Department of Immunology, University Medical Center Utrecht, The Netherlands; Department of Rheumatology & Clinical Immunology, University Medical Center Utrecht, The Netherlands

**Keywords:** keratoconus, GWAS, cellular aging, MTOR, RNA expression

## Abstract

**Purpose:** Keratoconus (KC) is an eye condition that can lead to a severe vision loss and may warrant a corneal grafting procedure. Meta-analyses of genome wide association studies have identified several genes that confer risks for differences in corneal curvature, corneal thickness, and developing keratoconus. Currently, there is limited evidence of a functional role for the identified loci in the affected corneal tissues.

**Methods:** We investigated the gene expression profiles of 4 GWAS confirmed risk loci and several related pathways that function in cellular ageing and cell cycle control in corneal tissue of a discovery and replication cohort comprising in total 27 keratoconus patients, 16 healthy controls, and 21 diseased controls (failed corneal grafts).

**Results:** We confirmed the *MTOR* gene locus as differentially expressed in KC corneas in a discovery cohort Next, we replicated these results in a second cohort and found evidence of increased expression of various mTORC1 pathway signature genes, namely *MTOR* itself (*P*=0.040), *AKT1* (*P*=0.028), *IGF1R* (*P*=0.022) and *RAPTOR* (*P*=0.007).

**Conclusions:** Gene expression profiling in cornea tissues revealed robust up-regulation of the mTORC1 pathway in KC and substantiates a potential role for this pathway in its pathogenesis. Functional implications should be further studied since biomarkers for disease activity are needed and selective targeting of the mTOR pathway is a promising treatment concept.

## Introduction

Keratoconus (KC) is an eye condition that leads to poor visual acuity due to myopia (short-sightedness) and irregular astigmatism, and in advanced cases to blindness due to corneal scarring. This is caused by the loss of corneal tissue, which precedes an archetypical cone-like shape of the cornea in KC.^1^ Sever cases warrant corneal transplant surgery to restore visual acuity. The underlying etiology is considered a complex interplay of corneal tissue remodeling, and activation of several proteases and inflammatory pathways.^4^ Interestingly, the high degree of concordance in monozygotic twins, and a high prevalence of KC in first degree relatives, indicates genetic predisposition as a predominant contribution to KC susceptiblity.^6–7^ Indeed, genome wide association studies (GWAS) have revealed susceptibility loci (*FOXO1* and *FNDC3B)* linked to central corneal thickness and keratoconus.^8^ Additionally, meta-analyses of large European and Asian cohorts reported that variants near *FNDC3B^9^, FRAP1/MTOR^9^*, and *PDGFRA^10^* genes conferred relatively large risks for corneal curvature aberrations- a hallmark of keratoconus. Unfortunately, the translation from genetic studies to functional understanding, let alone targeted therapeutic avenues, to combat corneal dysregulation in KC, is still in its infancy.

We hypothesized that the genetic changes put forward by GWAS in keratoconus and corneal curvature might highlight pathways that are actually deregulated in the cornea, affecting its architecture. To test this hypothesis we set out to map the expression of several associated risk genes in corneal tissues of patients to assess the actual implications of previous genetic studies *ex vivo*. We first screened a a discovery cohort (n = 36) for gene expression profiles of several genes and pathways highlighted by previous genetic studies in corneal tissue from KC patients, healthy controls, and diseased controls (decompensated corneal grafts). We technically validated the outcomes of the discovery cohort and used an independent replication cohort (n = 27) to assess the robustness and reproducibility of our observations.

## Methods

### Acquisition of corneal samples

KC cornea samples were collected from patients receiving a corneal transplant for severe KC (N=25) or pellucid marginal degeneration (N=1). Twenty-seven corneas from 27 patients were included in this group. The group of diseased controls are composed of decompensated grafts (DG). Twenty-one samples from 21 patients were included in this group. All aforementioned cornea buttons were processed using Tissue-Tek (Sakura Finetek U.S.A., Inc.) immediately after resection, cut into five full thickness slices, and stored at -80 °C. Healthy cornea (HC) controls were obtained from the Euro Cornea Bank (ECB), Beverwijk, The Netherlands, and Department of Anatomy, University Medical Center Utrecht, The Netherlands. A total of 16 corneas were prepared from post-mortem tissue within 24h of death and prepared from eyes from unrelated Caucasian donors who had no history of keratoconus, ocular inflammation, or vitreoretinal disease. The replication cohort comprised 11 KC, 11 DG, and 5 HC-samples, see figure 1.

**Figure 1.**
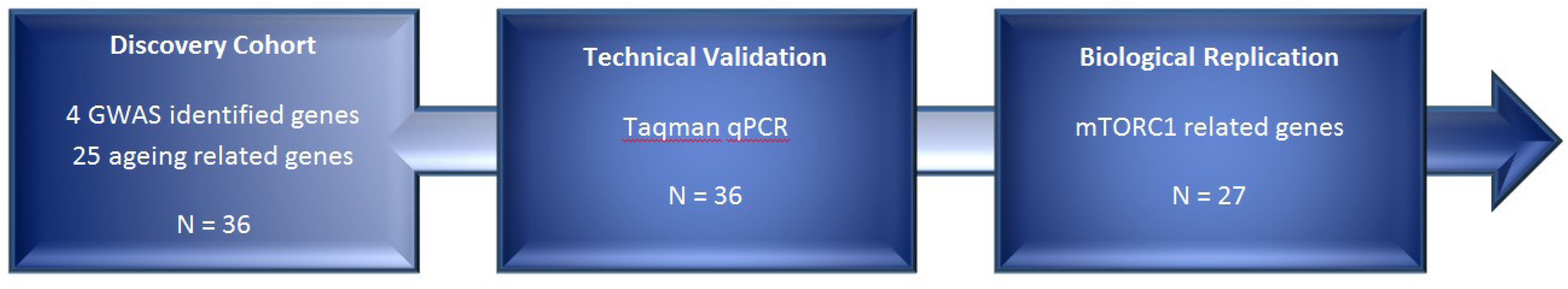
Flowchart of gene expression analyses and technical validation in discovery and replication cohort (N=64). Figure legend: GWAS = genome wide association study; mTORC1 = mammalian target of rapamycin 1 pathway.

This study was approved by The Medical Research Ethics Committee of the UMC Utrecht. It reviews research protocols in accordance with the Medical Research Involving Human Subjects Act (WMO). The MREC of the UMC Utrecht is accredited by the Central Committee on Research Involving Human Subjects (CCMO) since november 1999. The MREC of the UMC Utrecht is also member of the Dutch union of MRECs (NVMETC). None of the donors were from a vulnerable population and all donors or next of kin provided written informed consent that was freely given.

### Clinical data extraction

Patient records were reviewed for additional data collection, such as patient history and preoperative assessment, including slit lamp evaluation, Schirmer’s testing, and Scheimpflug corneal tomography (Pentacam HR, Oculus GmbH). Available data for the healthy control group was limited to age, sex and cause of death.

### RNA isolation

RNA isolation from corneal buttons was performed using the TRIzol method (Life Technologies, Thermo Fisher Scientific, USA) following the manufacturers protocol. RNA was isolated after dissolving the cornea in TRIzol reagent. After checking RNA integrity, complementary-DNA synthesis was performed using the iScript cDNA synthesis kit (Bio-Rad, USA).

### Gene expression analysis

Gene expression analysis was performed by OpenArray quantitative real-time polymerase chain reaction (qPCR), using *GUSB* and *GAPDH* as housekeeping genes for measuring relative expression levels. All qPCR analyses were performed on the QuantStudio 12K Flex Real-Time PCR System (Life Technologies, Thermo Fisher Scientific, USA). The qPCR data were interpreted using ExpressionSuite software version 1.0.3 (Life Technologies, Thermo Fisher Scientific, USA), The genes *AHRR, FOXO3, IL6, IL10* and *hTERT* did notreach detectable levels in our qPCR assay. Six healthy control samples, two keratoconus samples, and four decompensated grafts did not meet qPCR quality control criteria for RNA expression analysis, thus reducing the effective sample size from 64 to 51 corneas. Baseline characteristics of the non-viable KC and DG samples did not differ from the mean group.

### Data analysis

Gene expressions were presented as fold changes per sample and plotted in grouped scatter plots. Statistical analysis are reported threefold; firstly a comparison of marker levels between KC vs. HC, secondly a comparison of KC vs. both healthy and diseased control groups (HC+DG), and thirdly a comparison of HC vs. both diseased groups (KC+DG). Differences in marker levels were statistically tested using the one-way independent ANOVA for normal distributions or the Kruskall-Wallis Test for non-normal distributions. We used either Tukey's or Dunn’s Tests for Post Hoc multiple comparisons. Thresholds for significance were corrected for multiple testing, based on the number of genes per analysis (threshold for significance = 0.05/n). Multiple imputation was performed using a multivariable imputation method considering all entered variables. Statistical analyses were performed using SPSS 21.0 (IBM SPSS Statistics, USA) and graphs were made in Prism 6.02 (GraphPad Software Inc., USA).

## Results

### Study population

The mean age of the KC patients was 44.1±15.5 years, 66.9±12.2 years for the DG group, and 85.2±8.0 years for the healthy control group. Baseline characteristics (age, gender, atopy, contact lens use) of the replication cohort did not differ materially from the samples used in the discovery cohort. Demographics of the discovery and validation cohort are described in Table 1.

**Table 1.**
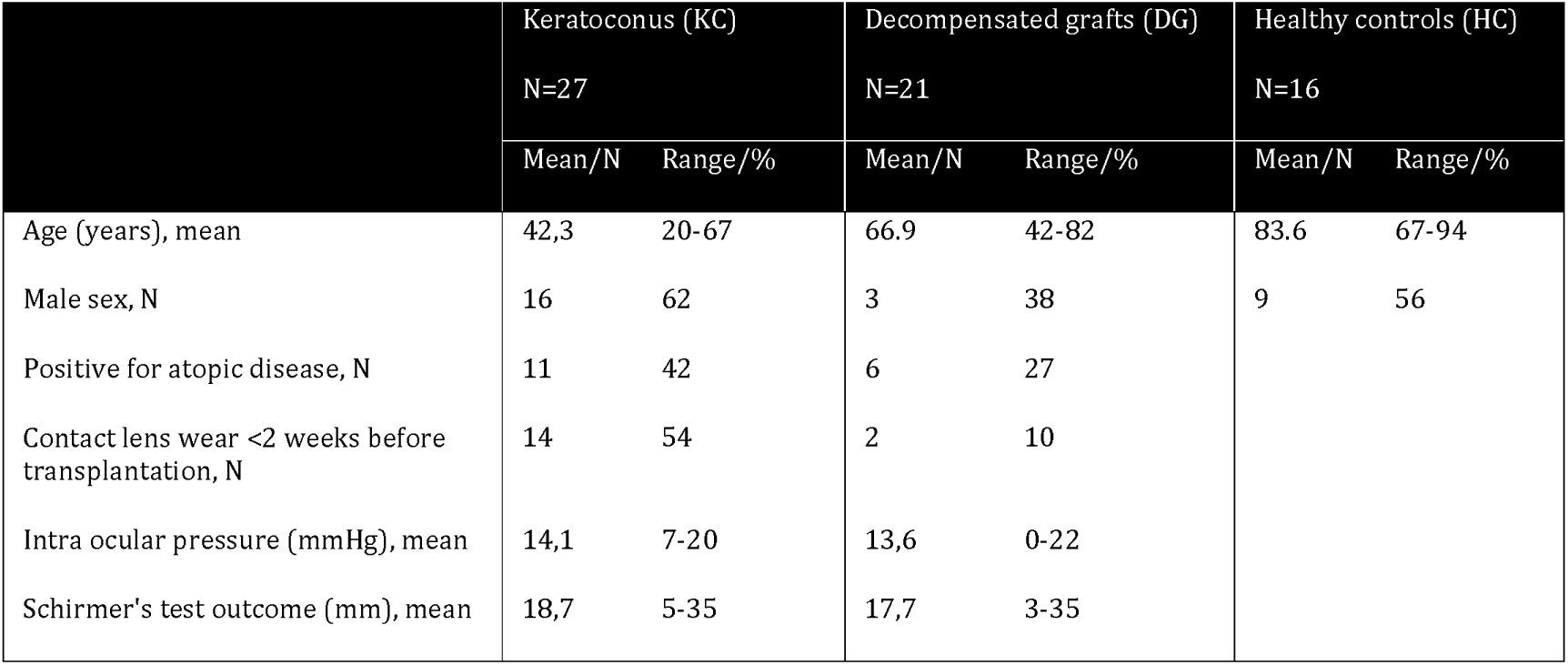
Study population characteristics.

|  | Keratoconus (KC) |  | Decompensated grafts (DG) |  | Healthy controls (HC) |  |
| --- | --- | --- | --- | --- | --- | --- |
|  | N=27 |  | N=21 |  | N=16 |  |
|  | Mean/N | Range/% | Mean/N | Range/% | Mean/N | Range/% |
| Age (years), mean | 42,3 | 20-67 | 66.9 | 42-82 | 83.6 | 67-94 |
| Male sex, N | 16 | 62 | 3 | 38 | 9 | 56 |
| Positive for atopic disease, N | 11 | 42 | 6 | 27 |  |  |
| Contact lens wear <2 weeks before transplantation, N | 14 | 54 | 2 | 10 |  |  |
| Intra ocular pressure (mmHg), mean | 14,1 | 7-20 | 13,6 | 0-22 |  |  |
| Schirmer's test outcome (mm), mean | 18,7 | 5-35 | 17,7 | 3-35 |  |  |

### Gene expression profile

Fifteen genes were significantly different expressed between KC vs. HC, and most of these genes were also affected in the diseased control group (DG). Of the four GWAS previously identified risk loci only *FRAP1/MTOR* showed a significantly altered expression in KC samples (*P*=0.005). Subsequently, several genes related to the mammalian target of rapamycin (mTORC1) pathway were significantly higher expressed in the KC samples compared to healthy controls: *AKT1* (24.8x higher, *P<*0.001), *DEPTOR* (4.9x, *P=*0.006), *FOXO4* (58.8x, *P<*0.001), *IGF1* (16.5x, *P<*0.001), *IGF1R* (20.4x, *P<*0.001), *MTOR* (6.5x, *P=*0.004), and *RAPTOR* (4.8x, *P=*0.010). In contrast, the levels of *MDM2* (0.16x, *P=*0.005) decreased. The expression of *RICTOR* (0.4, *P=*0.331) was not significantly altered. Strikingly, the aberrant gene expression profile of KC largely overlaps with severely failed corneal grafts (DG). Finally, the levels of *NFKB1* (13.9x, *P<*0.001), *SIRT7* (52.4x, *P<*0.001), and *WRN* (16.2x, *P<*0.001) were significantly lower in KC and DG when compared to healthy controls. Table 2 highlights the gene expression of the GWAS conformed loci, and mTORC1-pathway associated genes. Gene expression profiles of all genes of the discovery cohort are indicated in appendix 1, and visually represented in appendix 2 and 3.

**Table 2:**
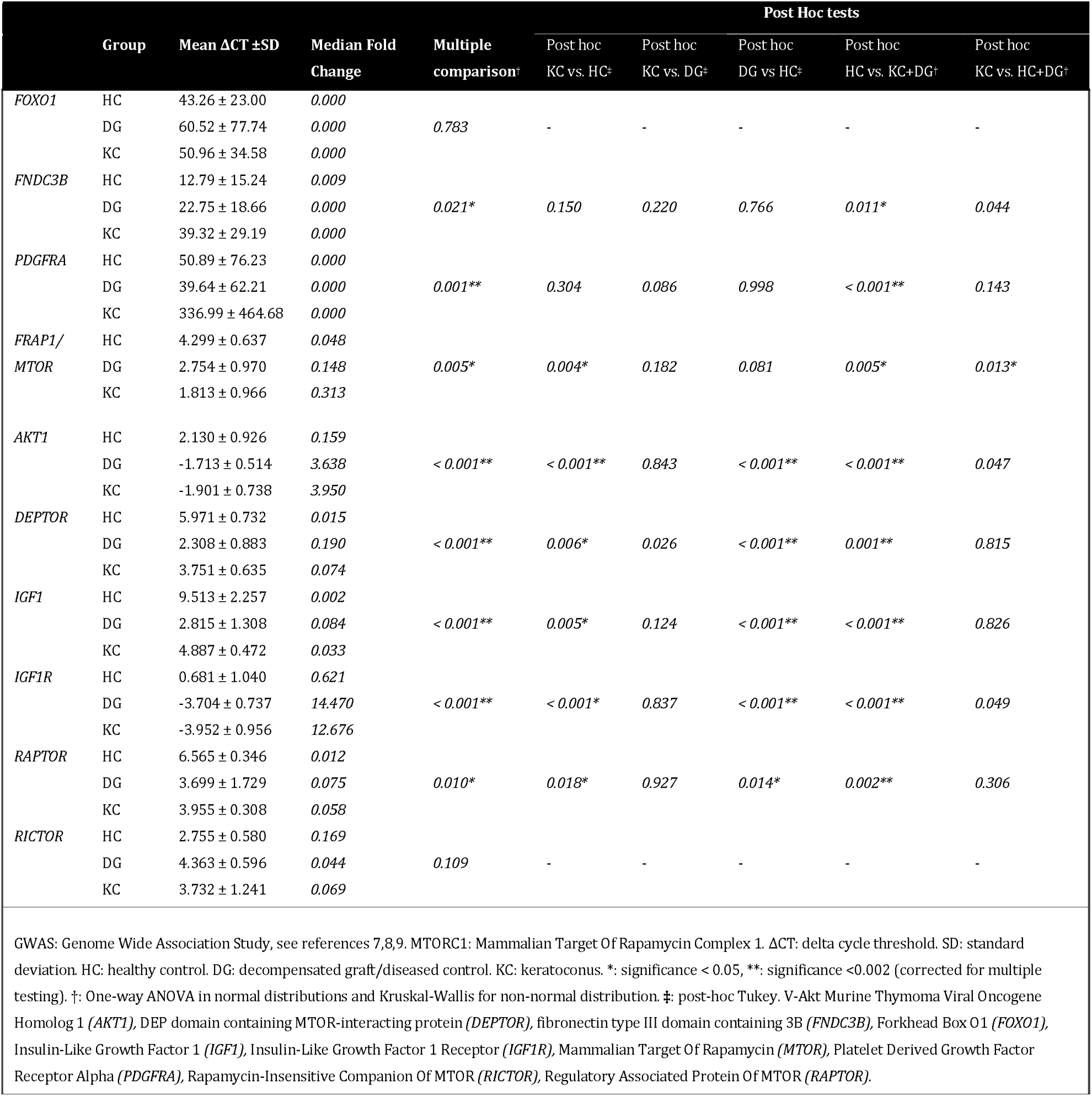
Gene expressions, median fold change and statistical analyses of GWAS identified genes and MTORC1-associated genes in the discoveiy cohort.

### Replication study

Since the mTOR gene was the only GWAS locus that displayed differential expression in corneas, we selected the mTORC1-pathway for replication analysis. In an independent cohort *AKT1, DEPTOR, IGF1, IGF1R, MTOR RAPTOR, REPTOR*, and *RICTOR* expression levels were assessed in 27 additional corneal samples (11 KC, 11 diseased controls and 5 healthy control samples). A solid difference in expression was noted for 4 out 7 genes involved in the mTORC1 pathway. A significantly altered gene-expression among all three sample groups in total was identified *for AKT1* (*P* = 0.028), *IGF1R* (*P* = 0.022), *MTOR (P* = 0.040), and *RAPTOR* (*P* = 0.007). Gene expression differences were less pronounced in the replication analysis, and mTORC1 pathway dysregulation was comparable between KC and diseased control samples, as was reported in the discovery cohort Details on post-hoc tests are given in Table 3.

**Table 3:** Gene expression of the mTORC1-pathway in keratoconus corneas, healthy controls and diseased controls in the pooled discovery and replication cohort.

| | Group | Mean $\Delta CT \pm SD$ | Median Fold Change | Multiple comparison<sup>a</sup> | Post Hoc tests | | | | |
| --- | --- | --- | --- | --- | --- | --- | --- | --- | --- |
|  |  |  |  |  | Post hoc KC vs. HC<sup>b</sup> | Post hoc KC vs. DG<sup>b</sup> | Post hoc DG vs HC<sup>b</sup> | Post hoc HC vs. KC+DG<sup>a</sup> | Post hoc KC vs. HC+DG<sup>a</sup> |
| *AKT1* | HC | 2.952±2.949 | 0.062 |  |  |  |  |  |  |
|  | DG | 5.100±2.852 | 0.031 | 0.028* | 0.110 | 0.699 | 0.024* | 0.010* | 0.616 |
|  | KC | 4.533±1.124 | 0.064 |  |  |  |  |  |  |
| *DEPTOR* | HC | 4.889±3.542 | 0.065 |  |  |  |  |  |  |
|  | DG | 7.109±3.471 | 0.009 | 0.074 | - | - | - | - | - |
|  | KC | 6.828±2.059 | 0.008 |  |  |  |  |  |  |
| *IGF1* | HC | 5.856±4.192 | 0.608 |  |  |  |  |  |  |
|  | DG | 7.457±3.649 | 0.654 | 0.345 | - | - | - | - | - |
|  | KC | 6.954±1.945 | 1.065 |  |  |  |  |  |  |
| *IGF1R* | HC | 0.785±2.975 | 0.239 |  |  |  |  |  |  |
|  | DG | 3.204±2.995 | 0.153 | 0.022* | 0.081 | 0.749 | 0.020* | 0.007** | 0.533 |
|  | KC | 2.646±1.808 | 0.241 |  |  |  |  |  |  |
| *MTOR* | HC | 4.610±3.082 | 0.013 |  |  |  |  |  |  |
|  | DG | 6.814±2.981 | 0.011 | 0.040* | 0.324 | 0.373 | 0.031* | 0.031* | 0.894 |
|  | KC | 5.861±2.608 | 0.023 |  |  |  |  |  |  |
| *RICTOR* | HC | 5.894±3.657 | 0.004 |  |  |  |  |  |  |
|  | DG | 7.967±3.187 | 0.003 | 0.098 | - | - | - | - | - |
|  | KC | 7.544±1.934 | 0.008 |  |  |  |  |  |  |
| *RPTOR* | HC | 2.952±2.949 | 0.052 |  |  |  |  |  |  |
|  | DG | 5.100±2.852 | 0.020 | 0.007** | 0.058 | 0.699 | 0.024* | 0.003** | 0.894 |
|  | KC | 4.533±1.124 | 0.047 |  |  |  |  |  |  |

ΔCT: delta cycle threshold. SD: standard deviation. HC: healthy control. DG: decompensated graft/diseased control. KC: keratoconus. \*: significance < 0.05, \*\*: significance < 0.008 (corrected for multiple testing). a: One-way ANOVA in normal distributions and Kruskal-Wallis for non-normal distribution. b: post-hoc Tukey. V-Akt Murine Thymoma Viral Oncogene Homolog 1 (*AKT1*), DEP domain containing MTOR-interacting protein (*DEPTOR*), Insulin-Like Growth Factor 1 (*IGF1*), Insulin-Like Growth Factor 1 Receptor (*IGF1R*), Mammalian Target Of Rapamycin (*MTOR*), Rapamycin-Insensitive Companion Of MTOR (*RICTOR*), Regulatory Associated Protein Of MTOR (*RPTOR*).

### Cluster analysis and visual representation of mTORC1 pathway expression

To reveal the underlying structure of mTORC1 pathway dysregulation, all corneal samples were subjected to unsupervised hierarchical clustering. Global comparisons by hierarchical cluster analysis discerned three overarching groups labeled C1, C2 & C3 (Figure 2). The three clusters roughly corresponded with each of the three investigated sampled specimens (keratoconus, decompensated grafts, healthy controls). C1 is the most homogenous set and contains largely healthy controls, whereas most keratoconus cases were found in C2 (17/25). Strikingly, keratoconus cases were also well represented in the C3 group, to a level comparable to the diseased controls. In the latter mTORC1 pathway dysregulation was most outspoken. A classification based on mTORC1 pathway activation was not readily feasible, though it should be noted that this clustering was performed only within the mTORC1 pathway.

**Figure 2.**
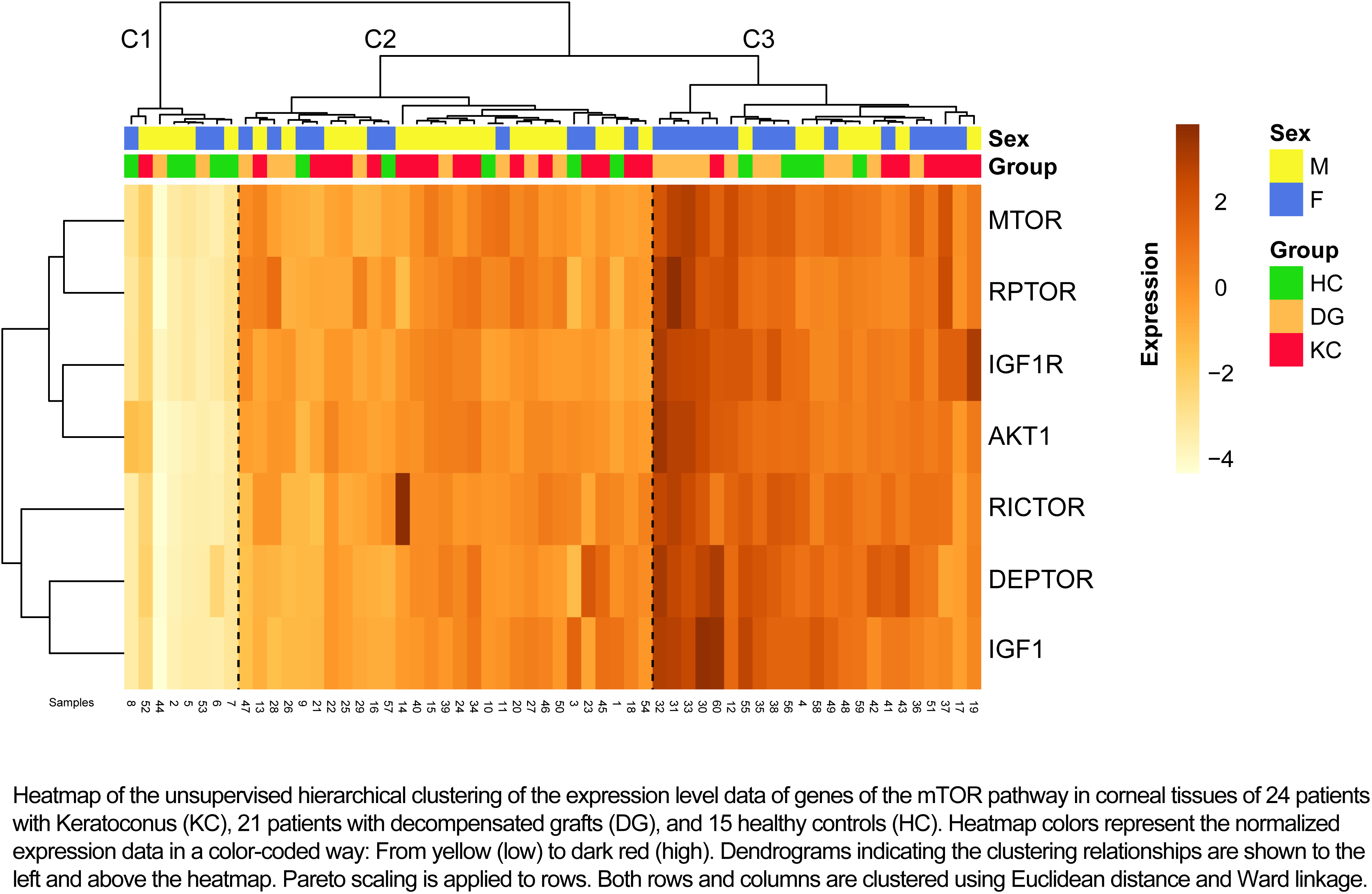
Heatmap of mTORC1 pathway gene expression in corneal samples of keratoconus patients, decompensated grafts, and healthy controls. Figure legend: Heatmap of the unsupervised hierarchical clustering of the expression level data of genes of the mTOR pathway in corneal tissues of 24 patients with Keratoconus (KC), 21 patients with decompensated grafts (DG), and 15 healthy controls (HC). Heatmap colors represent the normalized expression data in a color-coded way: From yellow (low) to dark red (high). Dendrograms indicating the clustering relationships are shown to the left and above the heatmap. Pareto scaling is applied to rows. Both rows and columns are clustered using Euclidean distance and Ward linkage.

## Discussion

In this study we confirm our hypothesis that susceptibility loci for keratoconus and corneal curvature point towards deregulation of pathways in the pathophysiology of keratoconus. Our results highlight especially the implication of the mTOR pathway. Insights provided by meta-analysis of GWAS data from large European and Asian KC cohorts have revealed susceptibility loci near *FOXO1* and *FNDC3B^8^*, and *MTOR/FRAP1^9^* and *PDGFRA^10^* in European and Asian cohorts that confer relatively large risk for the development of KC or biometric properties linked with KC (corneal thickness & corneal curvature). We only found the *mTOR* gene being differentially expressed in cornea’s from keratoconus patients compared to healthy controls. Interestingly, the levels of KC risk loci *FOXO1, FNDC3B* and *PDGFRA* were not significantly altered between the groups and suggest that the associated SNPs do not alter gene expression. Indeed, publicly available expression quantitative trait locus (eQTL) database Genevar revealed that the previously reported SNPs near *FOXO1, FNDC3B* or *PDGFRA* do not function in terms of transcript regulation of these genes.^8,11,12^

In addition to the *mTOR* gene itself, we identified several key components of the mTORC1 pathway to be significantly and reproducibly upregulated in KC corneas, including MTORC1 complex constituent *RAPTOR*, the gene coding the major growth factor *IGF1*, its receptor *IGF1R*, and *AKT1*, a potent stimulator of the *mTOR* pathway.^13^ These functional implications strengthen the previous genetic association with *MTOR/FRAP1* identified by genome-wide studies.

Although the absolute number of corneas investigated in this study is relatively small, flowing from the scarce availability of ex vivo corneas, we were able to reveal distinct gene expression profiles. Of note, the diseased corneal samples were all obtained during a grafting procedure of more severe keratoconus patients. Therefore our results are mainly applicable to more severe KC. To reduce technical bias all samples were handled and analyzed in the same laboratory utilizing the same protocol and apparatus.

Previous studies indicated a role for *mTOR* signaling in maintaining the corneal homeostasis. For instance, corneal wound healing in murine corneas seems to be mediated through *PDGF-BB* in a *mTOR* dependent matter.^14^ Since decompensated grafts exhibit an ongoing inflammatory process, this might be an explanation for the increased *mTOR* signaling observed in decompensated graft corneas compared to healthy controls. In addition, corneal scarring is common in the severe cases of keratoconus, while less advanced cases can also show the archetypical conical shape, though with a completely clear cornea. Importantly, the sample set in this study also included clear corneas and revealed mTORC pathway activation for these corneas as well. This might indicate that the increased mTOR signaling precedes the scarring and therefore might underlie the progressive tissue loss associated with advanced keratoconus. In this mouse model, the process of ongoing wound-repair responses was successfully inhibited by administering rapamycin intra-ocularly.^14^ In addition, *in vivo* and *in vitro* mTOR pathway inhibition has showed less corneal TGF-ß and myofibroblast activation^15^. Increased TGF-b signaling has been shown to be an important part of KC pathogenesis.^16^

The combination of mTOR signaling in the corneal wound repair and the implication of this pathway found in GWAS and our translational study, makes it tempting to speculate that the mTOR pathway opens novel diagnostic and therapeutic avenues in keratoconus.^17^ We believe that a future prospective clinical study could manifest mTOR as a biomarker for disease activity and its inhibition as an alternative for invasive corneal surgery.

## Acknowledgments

Annemieke Haasnoot, Fleurike Verhagen, Jan Beekhuis are thanked for their technical assistance, and Prof. Dr. Ronald Bleys, Dr. Annette Gijsbers-Bruggink and Dr. Pieter Jan Kruit for the procurement of the corneal samples.

**Abbreviations**
ACT: delta cycle threshold
AHR: Aryl Hydrocarbon Receptor gene
AHRR: Aryl-Hydrocarbon Receptor Repressor gene
AKT1: V-Akt Murine Thymoma Viral Oncogene Homolog 1 gene
CDKN2A: Cyclin-Dependent Kinase Inhibitor 2A gene
CT: cycle threshold
DEPTOR: DEP domain containing MTOR-interacting protein
DG: decompensated graft
ECB: European Cornea Bank, Beverwijk, The Netherlands
FNDC3B: fibronectin type III domain containing 3B
FOXO1: Forkhead Box O1 gene
FOXO3: Forkhead Box O3 gene
FOXO4: Forkhead Box O4 gene
FRAP1: MTOR = Mechanistic Target Of Rapamycin gene
GWAS: Genome Wide Association Study
H2AFX: H2A Histone Family, Member X gene
HC: healthy control
HDAC9: Histone Deacetylase 9 gene
IGF1: Insulin-Like Growth Factor 1 gene
IGF1R: Insulin-Like Growth Factor 1 Receptor gene
IL6: Interleukin 6 gene
IL10: Interleukin 10 gene
KC: Keratoconus
MDM2: Mouse Double Minute 2 homolog gene
MTOR: FRAP1 = Mechanistic Target Of Rapamycin gene
mTOR: mammalian target of rapamycin protein
mTORc1: mammalian target of rapamycin complex 1
mTORC2: mammalian target of rapamycin complex 2
NFATC1: Nuclear Factor of Activated T-cells, Cytoplasmic 1 gene
NFKB1: Nuclear Factor Of Kappa Light Polypeptide Gene Enhancer In B-Cells 1 gene
PDGFRA: Platelet Derived Growth Factor Receptor Alpha
PTEN: Phosphatase And Tensin Homolog gene
qPCR: quantitative real-time polymerase chain reaction
RICTOR: Rapamycin-Insensitive Companion Of MTOR gene
RAPTOR: Regulatory Associated Protein Of MTOR gene
SIRT1: Sirtuin 1 gene
SIRT6: Sirtuin 6 gene
SIRT7: Sirtuin 7 gene
TERT: Telomerase Reverse Transcriptase gene
TP53: Tumor Protein P53 gene
WRN: Werner syndrome, RecQ helicase-like, gene

